# UMIche: A platform for robust UMI-centric simulation and analysis in bulk and single-cell sequencing

**DOI:** 10.1101/2025.03.15.643072

**Authors:** Jianfeng Sun, Shuang Li, Stefan Canzar, Adam P. Cribbs

## Abstract

Unique molecular identifiers (UMIs) have actively been utilised by various RNA sequencing (RNA-seq) protocols and technologies to remove polymerase chain reaction (PCR) duplicates, thus enhancing counting accuracy. However, errors during sequencing processes often compromise the precision of UMI-assisted quantification. To overcome this, various computational methods have been proposed for UMI error correction. Despite these advancements, the absence of a unified benchmarking and validation framework for UMI deduplication methods hinders the systematic evaluation and optimisation of these methods. Here, we present UMIche, an open-source, UMI-centric computational platform designed to improve molecular quantification by providing a systematic, integrative, and extensible framework for UMI analysis. Additionally, it supports the development of more effective UMI deduplication strategies. We show that through its integration of a broad spectrum of UMI deduplication methods and computational workflows, UMIche significantly advances the accuracy of molecular quantification and facilitates the generation of high-fidelity gene expression profiles.

## Introduction

RNA sequencing (RNA-seq) technologies have been advanced through optimised sequencing protocols that enable the generation of high-quality transcriptional expression profiles [1–3]. Central to these optimisations are the accurate sequencing of nucleotides [4] and the effective quantification of reads [5], which support both sequence-based [6,7] and count matrix-based studies [8,9]. Accurate molecular quantification has garnered increasing attention due to its critical role in elucidating gene regulation mechanisms across diverse cellular functions [10,11]. To achieve precise gene expression quantification, the attachment of Unique Molecular Identifiers (UMIs) to the sequencing molecule at the 3’ or 5’ end has been widely adopted [12]removes PCR duplicates in the majority of sequencing protocols [13–15]. The use of UMIs maximises counting accuracy while minimising complexity, particularly in cDNA libraries [13]. However, the error-prone nature of PCR amplification and sequencing processes can reduce the accuracy of UMI-based quantification [16,17]. Therefore, numerous computational methods and tools have been developed for UMI error correction [18–21].

Current strategies for UMI collapsing or deduplication predominantly rely on edit distance-dependent methods [22–26]. Notable examples, including UMI-tools [27] and UMICollapse [28], which create UMI graphs with an edit distance threshold to merge compositionally similar UMIs, assuming them to originate from the same molecule. Conversely, a few methods, such as DAUMI [29] and TRUmiCount [30], employ alternative approaches that model and evaluate the distributions or abundances of potential PCR duplicates to effectively collapse UMIs.

Despite these advancements, there remains a need for an integrated platform that supports systematic analysis, evaluates established methods, and fosters the development of new ones. The recent introduction of homoblock UMI correction approaches has allowed the leveraging of recurrence patterns in homoblocks for error correction [31,32]. Consequently, new error correction strategies, such as majority voting and set cover optimisation, have been developed [33,34]. These strategies enable the collapsing of nucleotides within dimer and trimer blocks (termed intra-molecular collapsing) before collapsing between UMIs (termed inter-molecular collapsing). However, there is a lack of systematic benchmarking for these studies.

Here, we introduce a **UMI** tool ni**che** (UMIche), a comprehensive platform that integrates multiple UMI collapsing methods and streamlines computational pipelines to facilitate the evaluation and development of strategies for removing PCR duplicates. UMIche operates at the platform level, harmonising UMI approaches from several different sequencing protocols, calculating deduplication counts, exploring features of different methods and visualising analysis results. To better support the UMI-centric analysis, we further provide a deep learning-based framework for on-demand simulation of UMI count matrices at the single-cell level. These data can then be used to generate sequencing reads to aid in benchmarking UMI collapsing tools. Overall, UMIche aims to enhance understanding of UMI collapsing by optimising the combination and use of existing methods and data resources. We also demonstrate several use cases to illustrate how UMIche can be leveraged to achieve optimal performance in UMI deduplication.

## Results

### Overview of UMIche

UMIche is a UMI-centric platform designed to support the development and evaluation of UMI collapsing methods, using both external and internal simulated or experimental data across multiple sequencing contexts, such as bulk and single-cell levels. It is highly modular, featuring Python interfaces to facilitate seamless integration into other computational programs and analysis pipelines. UMIche comprises main four modules: UMI collapsing method implementation, UMI collapsing pipeline implementation, UMI count matrix implementation, and visualisation of various UMI metrics (**Figure 1**). The core of UMIche offers multifaceted functionalities for UMI deduplication at the single locus, bulk, single-cell levels.

**Figure 1.**
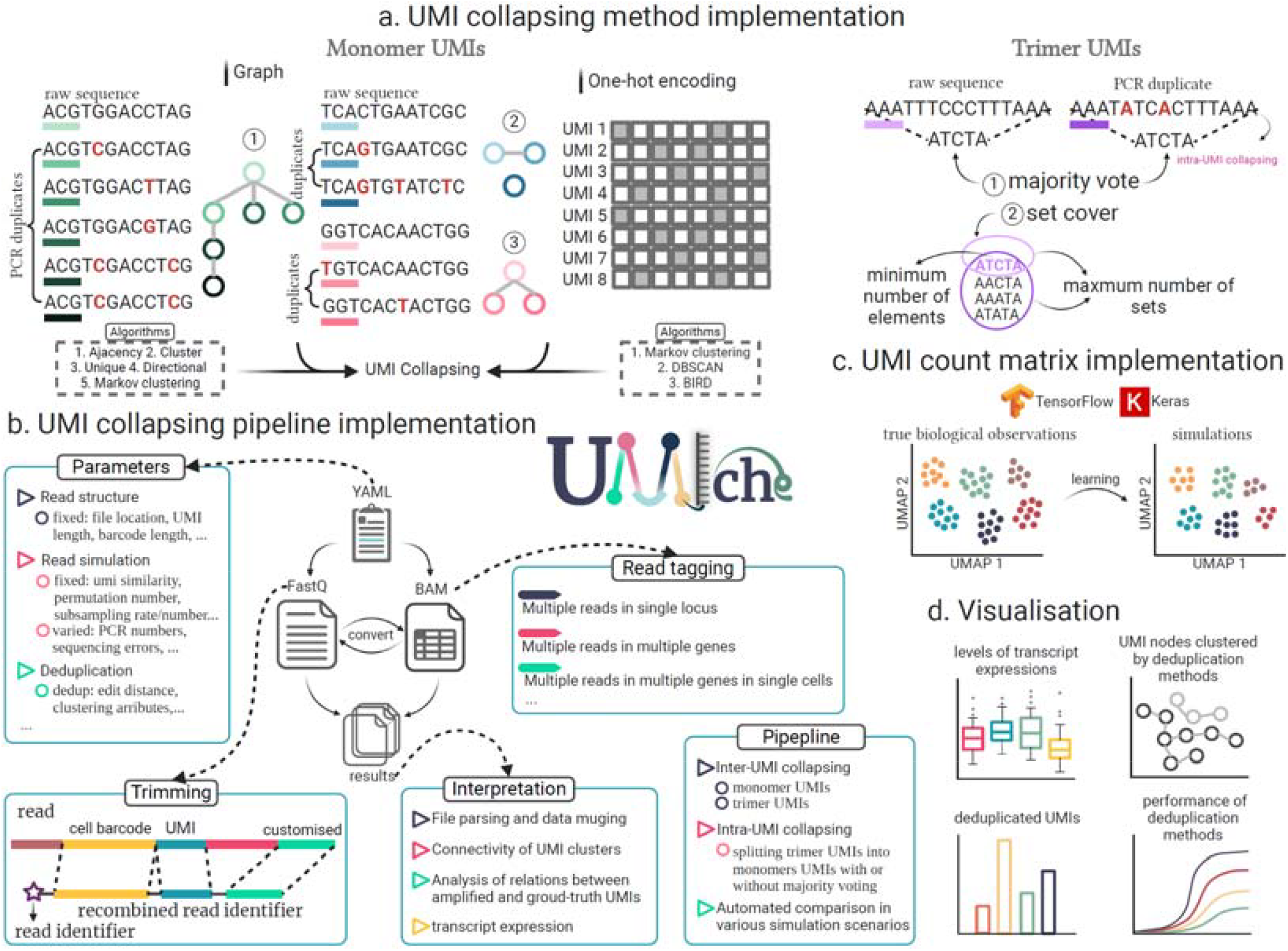
Overview of UMIche. It contains UMI collapsing method implementation (**a**), UMI collapsing pipeline implementation (**b**), UMI count matrix implementation (**c**), and visualisation of UMI deduplication metrics (**d**). All methods in mclUMI and UMI-tools have been re-implemented for deduplicating monomer UMIs. Majority voting and set coverage optimisation specialised for deduplicating UMIs of homoblocks have been made available. Together with 3 Euclidean distance-based clustering methods that process the one-hot encodings of UMIs sequences, there are a total of 12 UMI deduplication methods available in UMIche. We provide a module to simulate single-cell UMI count matrix on-demand with ground-truth labels, facilitating the examination of the performance of UMI deduplication methods. We built a few computational pipelines following the same protocol to pre- and post-process sequencing data, including read tagging, trimming UMIs from reads, collapsing UMIs, etc. This feature allows a rapid benchmark of performance of UMI deduplication tools on large-scale data. A plotting module is provided to readily visualise the performance of multiple UMI deduplication tools.

1. **UMI collapsing modules**. We include modules to handle the deduplication of PCR artefacts tagged with two types of UMIs inclusive of monomer and trimer UMIs (**Figure 1a**). Recently, the range of methods for UMI collapsing has broadened from traditional edit distance-dependent approaches (*e*.*g*., constructing UMI graphs with edit distance to assist molecular quantification) to methods that do not rely on edit distance, especially following the introduction of homoblock UMIs [31,32] (*e*.*g*., the majority vote and set cover optimisation methods). To comprehensively support UMI collapsing, UMIche includes executables for both approaches. The investigation into edit distance-free methods has enhanced our understanding of how closely the number of UMI-tagged reads can approximate the original transcript count under extreme conditions. UMIche thus facilitates the exploration of the limits of UMI collapsing methodologies.
2. **UMI collapsing pipeline**. To enable systematic and rapid analysis, UMIche provides workflows running at the pipeline level (**Figure 1b**). To harmonise experimental and simulated sequencing protocols, UMIche standardises input data formats by accepting read alignments in BAM file format [35] or converting FastQ reads to BAM files. To minimise the workload within Python scripts, we implemented a YMAL-dependent [36] analytical workflow, streamlining the initialisation of parameters required for both upstream and downstream analyses.
3. **Simulated ground-truth count matrix**. For single-cell studies, generating count matrices with ground-truth labels is essential for simulating reads, which are subsequently used for in silico testing of UMI collapsing methods or sequencing protocols (**Figure 1c**). To facilitate this, we implemented a deep learning-based framework, DeepConvCVAE, to provide users with count matrices on-demand. With a GPU support, this framework operates at high speed, efficiently handling the substantial data load of single-cell datasets. Users can retrain the deep learning model using their own data or generate a count matrix based on a built-in model for general testing purpose. Detailed descriptions of this framework are provided in the subsequent sections.
4. **Visualisation**. This module is designed to generate bar and line plots to facilitate the interpretation of analysis results (**Figure 1d**). It visualises deduplicated molecule counts and gene expression levels measured by different UMI collapsing methods. Additionally, we offer a graph plotting service to visually represent different UMI clusters. This feature aids in understanding how various UMIs are depicted in a UMI graph and partitioned into distinct clusters. Such visualisations can helps study the trajectories of tagging molecules (e.g., UMIs) across multiple PCR cycles and provide insights into how errors in UMIs evolve under varying conditions, such as increasing sequencing depths, higher down-sampling, and extending PCR cycle.

### Streamlining analysis with pipelines in UMIche

In UMIche, we have organised functionalities into specific pipelines to streamline the analysis process. There are two main pipelines: one dedicated to deduplicating UMIs (Step 3 in **Figure 2a**) and the other focused on UMIs trajectories (Step 4 in **Figure 2a**). The deduplication pipeline provides standard deduplication results, whereas the trajectory pipeline offers insights that enhance methodologies and improve the performance of UMI deduplication methods. Both pipelines are designed to be managed uniformly, with consistent parameter configuration and file reformatting, to minimise user effort.

**Figure 2.**
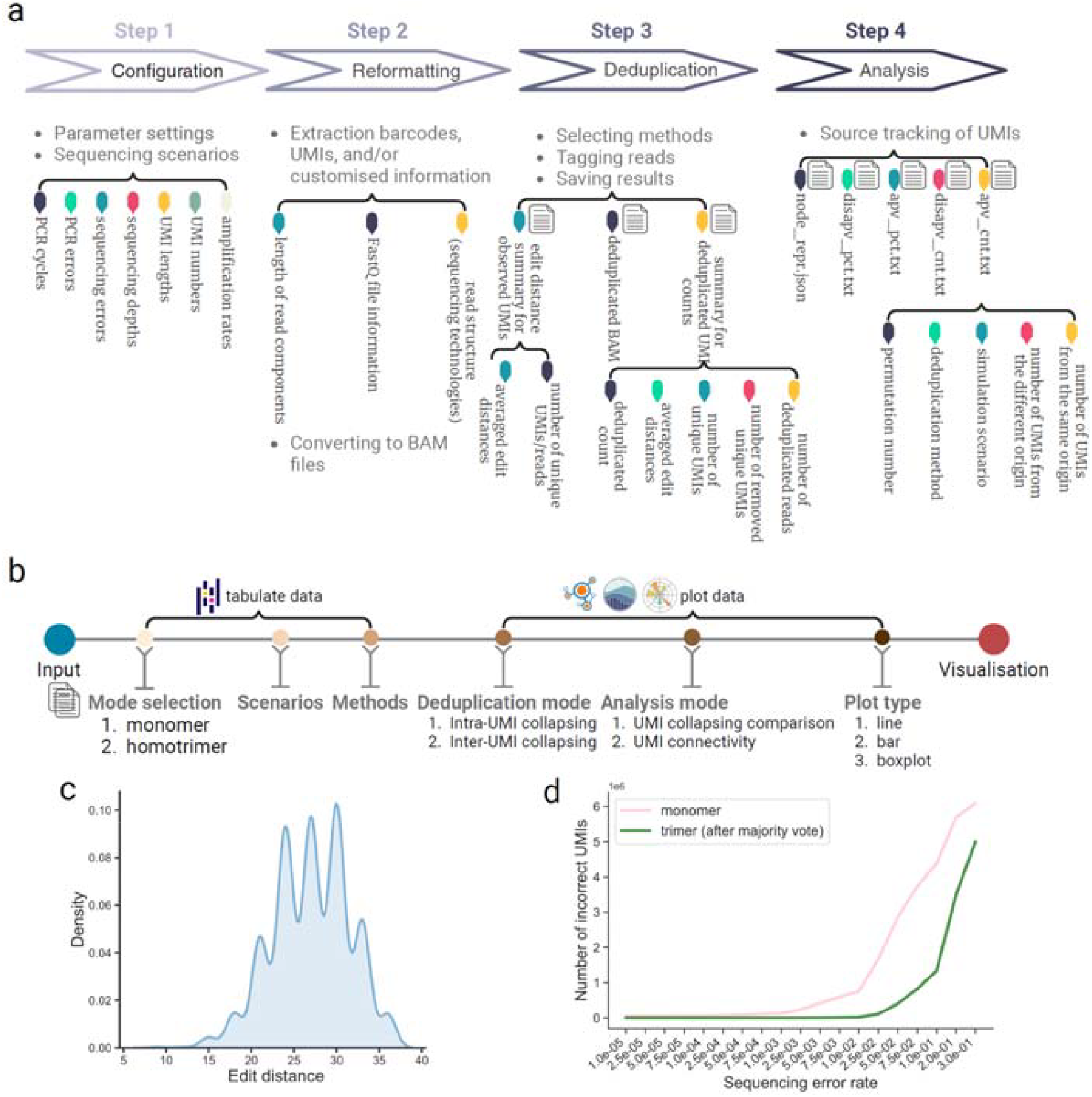
Workflow of the pipeline module in UMIche. **a**, typical modules in a UMIche pipeline. **b**, processing and visualisation of deduplication results. **c**, an example of visualising edit distances between simulated UMIs using a UMIche pipeline. **d**, an example of counting numbers of error monomer and trimer UMIs using a UMIche pipeline.

We propose strategies for both inter-molecular collapsing (collapsing between different UMIs) and also intra-molecular collapsing (collapsing of nucleotides within UMIs). Inter-molecular collapsing is used to eliminate PCR artefacts by collapsing UMIs, while homotrimer UMIs facilitate intra-molecular collapsing when redundant signals are detected between nucleotides in trimer blocks. Interestingly, intra-molecular collapsing can be incorporated at multiple stages of the inter-molecular collapsing process, maximising the performance gains in UMI deduplication results and providing new insight into UMI collapsing strategies.

The pipelines in UMIche automate these processes, facilitating the discovery of new knowledge using both simulated and experimental short-read and long-read data in bulk RNA-seq and scRNA-seq conditions. In the following sections, we present several use cases to illustrate how UMIche pipelines advance UMI deduplication performance.

### Case 1: Improving molecular deduplication computational strategies using homotrimer UMIs

Our efforts to accurately measure transcriptional expression profiles have led to the development of homoblock UMIs, which surpass the limitations of current computational demultiplexing approaches. This novel UMI deduplication method combines experimentally identifiable errors within the UMI sequence with computation-based sample demultiplexing, thereby enhancing UMI duplication. The introduction of homotrimer UMIs has enabled the use of a “majority vote” strategy and the implementation of a set cover optimisation technique, thanks to recurring nucleotides patterns. Despite their potential, a thorough comparative analysis of these approaches is needed to identify the most efficient method for specific sequencing contexts and to inform the development of future improvements.

To address this, we first review the process of removing PCR duplicates using homotrimer UMIs with majority vote and the set cover optimisation (**Figure 3**). The majority vote demultiplexing process involves two steps: (*i*) obtaining monomer UMIs by collapsing blocks of homotrimer UMIs (which we refer to as intra-molecular collapsing) and (*ii*) removing homotrimer UMIs according to repeated monomer UMIs (which we refer to as inter-molecular collapsing) (**Figure 3a and 3c**). The set cover demultiplexing process involves three steps: (*i*) splitting homotrimer UMIs into monomer UMIs, (*ii*) covering highly similar homotrimer UMI with their split monomer UMIs, and (*iii*) removing homotrimer UMIs according to repeated monomer UMIs (**Figure 3a and 3b**). It remains unclear whether further improvements in UMI deduplication performance can be achieved by deduplicating those retained homotrimer UMIs. As shown in **Figure 3d**, we are interested in utilising UMIche to perform the second round of intra-molecular collapsing of homotrimer UMIs and apply external graph-based UMI collapsing methods, built on top of the retained homotrimer UMIs.

**Figure 3.**
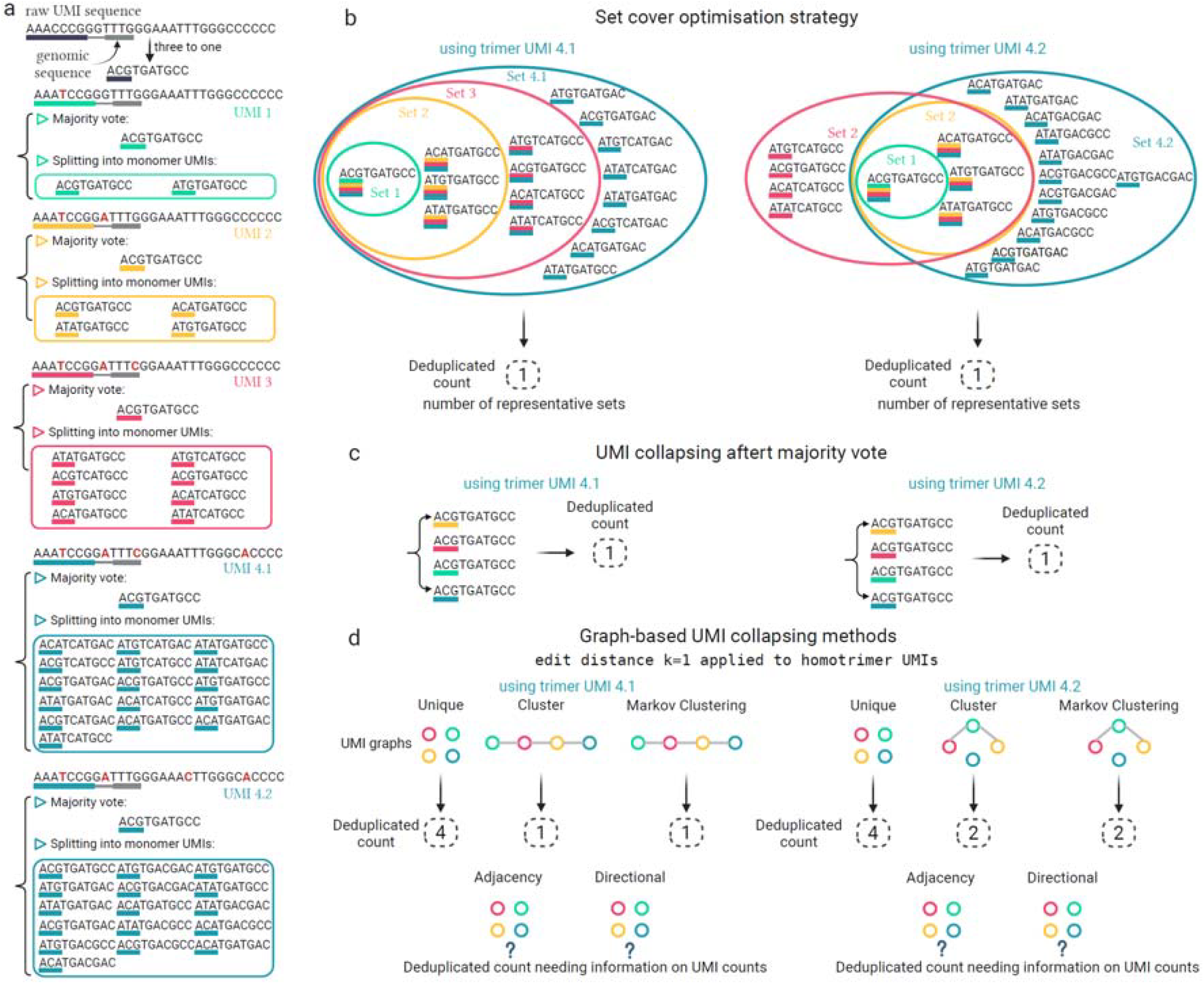
Working principle of the set cover optimisation strategy. **a**, an original homotrimer UMI and 5 its PCR duplicates with errors. **b**, covering monomer UMIs split from the homotrimer UMIs using the set cover optimisation technique. **c**, illustration of deduplicating the homotrimer UMIs from **a** using the majority vote strategy. **d**, illustration of deduplicating the homotrimer UMIs from **a** using graph-based UMI collapsing methods.

**Figure 4.**
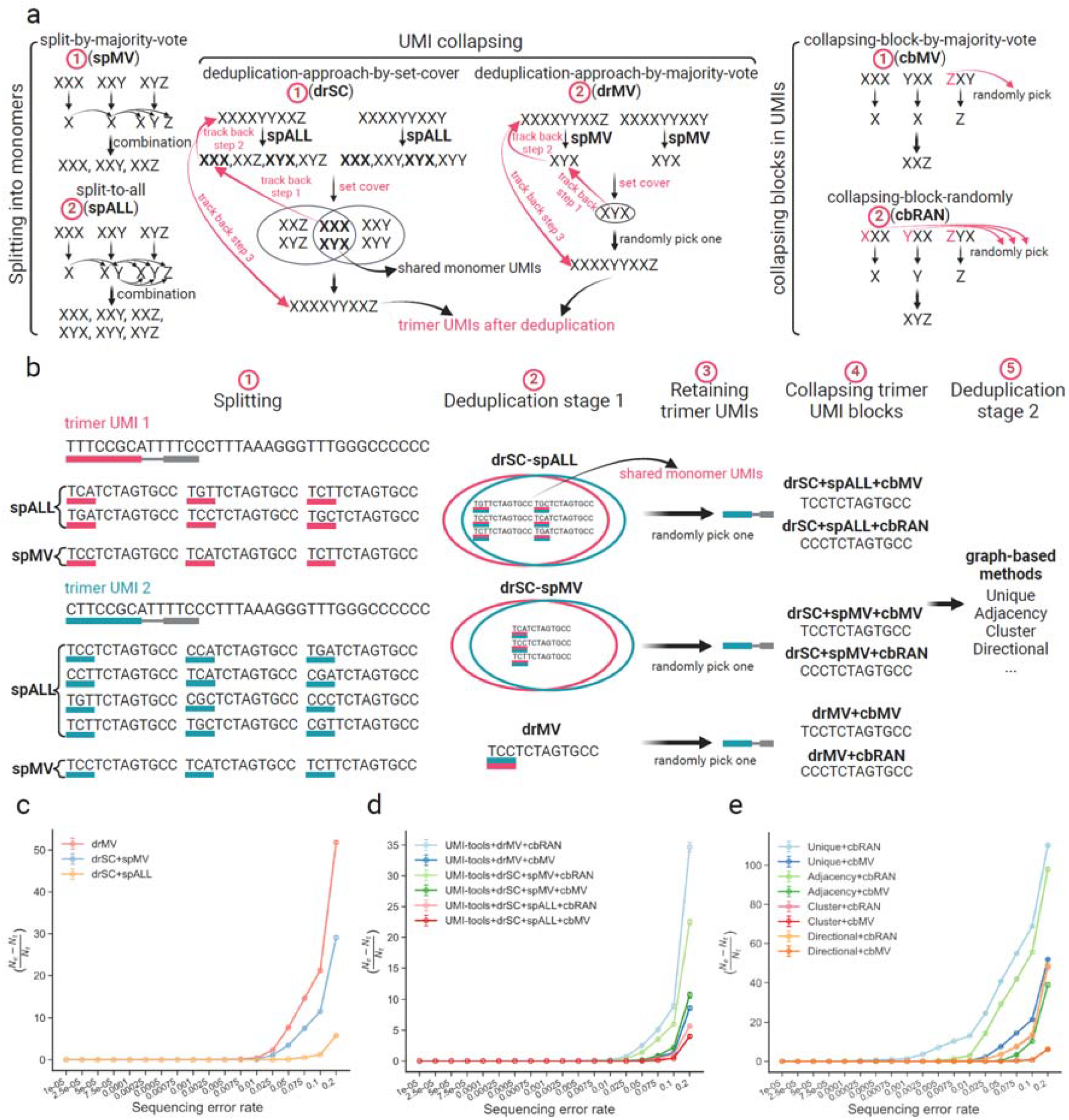
Comprehensive comparison between UMI collapsing methods. **a**, block-wise splitting homotrimer UMIs into monomer UMIs using the split-by-majority-vote (*spMV*) and split-to-all (*spALL*) strategies. Illustration of UMI collapsing by the set cover optimisation approach (*drSC*) and the majority vote approach (*drMV*). Illustration of collapsing blocks in UMIs by the majority vote strategy (*cbMV*) and the random selection (*cbRAN*). **b**, UMI collapsing by 1) splitting homotrimer UMIs into monomer UMIs, the first round of UMI collapsing, retaining homotrimer UMIs, collapsing UMI blocks, and the second round of UMI collapsing. **c**, Comparison of deduplication results between the *drMV* and *drSC* methods. *N*_*e*_ represents the estimated deduplicated count and *N*_*t*_ represents the true count. **d**, Comparison of UMI-tools deduplication results between the *drMV* and *drSC* methods. **e**, Comparison of deduplication results between different UMI-tools methods (*Unique, Adjacency, Cluster*, and *Directional*).

### 1. Boosting UMI deduplication effects by altering input materials to set cover optimisation

We first investigated whether alterations in the content of monomer UMIs input to set cover optimisation could lead to different molecular counting results. During the process of splitting homotrimer UMIs into monomer UMIs, we compared UMI deduplication performance using two strategies. The first strategy, *spMV*, involves splitting monomer UMIs from homotrimer UMIs by resolving nucleotide heterogeneity obviated in each trimer block using majority vote (**Figure 3a**). The second strategy, *spALL*, splits monomer UMIs without addressing nucleotide heterogeneity conflicts in trimer blocks. Our findings indicate that using set cover optimisation, the UMI deduplication performance with the *spALL* strategy outperforms that with the *spMV* strategy across a range of error rates (**Figure 3c**). Notably, the *spALL* strategy closely approximates the actual number of molecules, especially under conditions of high error rates. This makes the *spALL* strategy particularly useful for error-prone sequencing technologies or challenging sequencing conditions.

### 2. Comparison between majority voting and set cover optimisation

Next, we investigated whether the homotrimer UMIs resulting from the first-stage deduplication process could be further deduplicated in the second-stage process using external computational tools (e.g., UMI-tools). to select the most common nucleotide in each trimer block, with a random nucleotide chosen. To achieve this, we applied intra-molecular collapsing, combining three nucleotides in trimer blocks into monomer UMIs, as input for the well-established graph-based methods in UMI-tools (**Figure 3a and 3b**). The majority vote strategy was used if all three were different. This intra-collapsing strategy, termed *cbMV*, was used for the second round of UMI deduplication. We also developed another strategy, *cbRAN*, where a nucleotide is randomly selected from each trimer block without majority voting. Our results demonstrate that incorporating external computational tools significantly improves counting accuracy (**Figure 3c and 3d**), with the *cbMV* strategy yielding better results than *cbRAN*. This improvement is attributed to the inability of random nucleotide selection to correct errors effectively, particularly under high sequencing error rates. Our data suggests that majority voting can be progressively applied to enhance UMI deduplication across multiple stages. A comparison of methods within UMI-tools indicates that the Directional method remains the most accurate for molecular counting (**Figure 3e**), consistent with the main finding from previous studies [27].

### 3. Understanding the influence of monomer UMI splitting strategies on UMI deduplication effects

We then investigated how different UMI splitting strategies influence the performance of UMI deduplication. We calculated the accumulated number of homotrimer or monomer UMIs resolved or not resolved by the set cover optimisation technique. A group of homotrimer UMIs is considered resolved during the deduplication process if a monomer UMI, identified by the greedy algorithm as having the maximum count (see section *Methods* for greater details), is one of the UMIs collapsed from these homotrimer UMIs (**Figure 5a**). Otherwise, this UMI is counted as not resolved.

**Figure 5.**
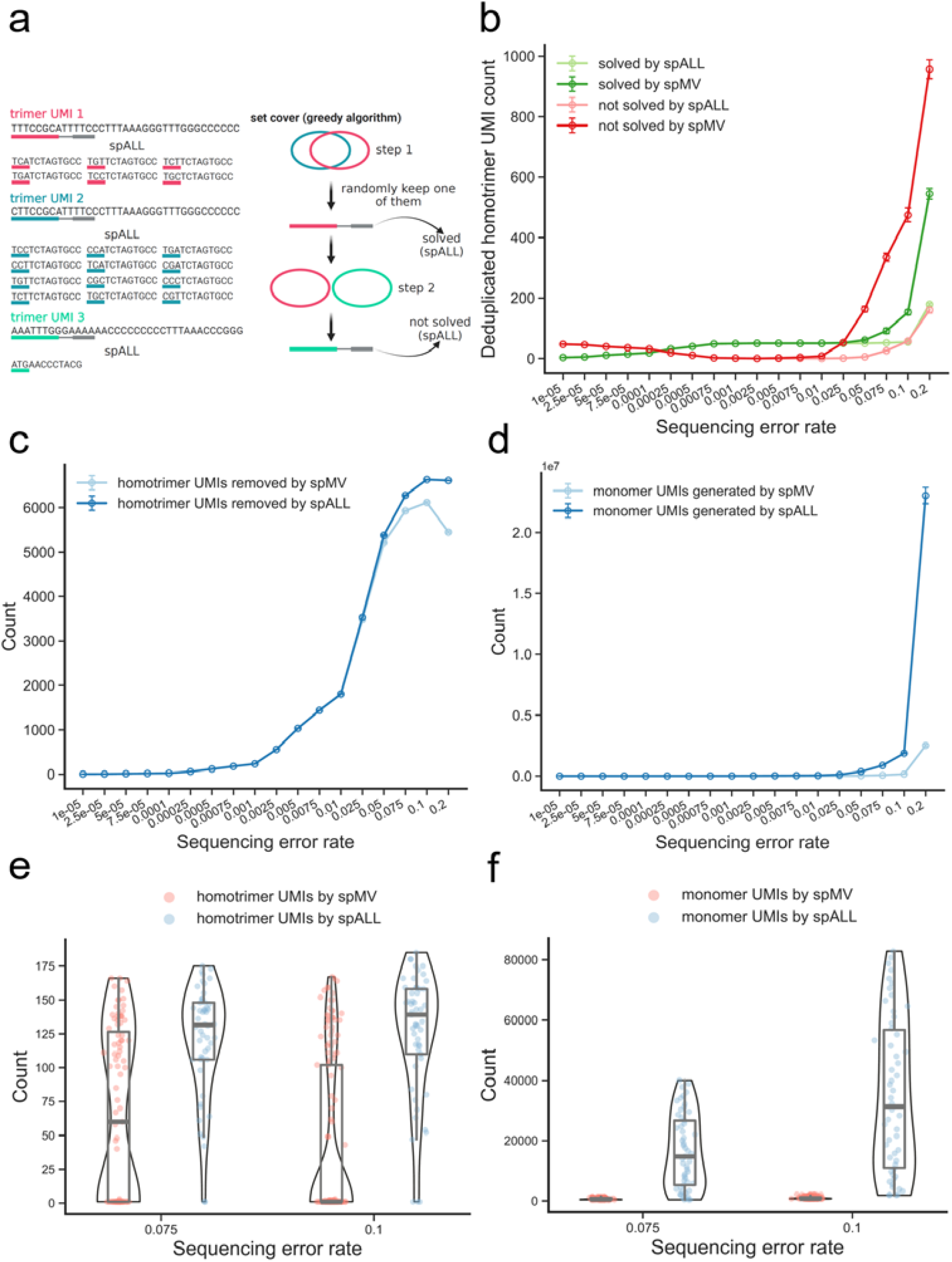
Analysis of split monomer UMIs and deduplicated homotrimer UMIs during set cover optimisation. **a**, illustration of solved UMIs by the set cover optimisation technique. **b**, Comparison of deduplicated counts of homotrimer UMIs solved by the set cover optimisation technique if the UMIs are split by *spMV* and *spALL*. **c**, Counts of homotrimer UMIs removed by set cover optimisation when the UMIs are split by *spMV* and *spALL*. **d**, Counts of monomer UMIs generated by *spMV* and *spALL* **e**, Counts of homotrimer UMIs removed per step using set cover optimisation when the UMIs are split by *spMV* and *spALL*. Each dot represents the count of removed homotrimer UMIs in each step of using the greedy algorithm for solving the set cover problem. **f**, Counts of monomer UMIs generated by *spMV* and *spALL* per step using set cover optimisation.

Our findings indicate that using UMIs split by the *spMV* strategy results in a similar deduplicated count to that obtained by the *spALL* strategy at low and moderate error rates. However, at high error rates, the *spMV* strategy leads to a significant increase in the deduplicated count (**Figure 5b**), accompanied by a sharp decrease in the number of homotrimer UMIs detected by the set cover optimisation for removal (**Figure 5c**). The final deduplicated count (or the accumulated number of removed homotrimer UMIs) is proportionally correlated with the number of the input monomer UMIs split by *spMV* and *spALL* (**Figure 5d**). Specifically, a higher number of input monomer UMIs correspond to more homotrimer UMIs, which are likely a result of PCR amplification (**Figure 5e and 5f**). One possible explanation is that the monomer UMIs split by *spMV* constitute a subset of all possible monomer UMIs split by *spALL*, making them less representative of homotrimer UMIs. This subset nature of monomer UMIs split by *spMV* reduces the likelihood of covering homotrimer UMIs effectively, thereby impacting the overall deduplication process.

### Case 2: DeepConvCVAE for simulating cell type-specific gene expression matrix for modelling UMI counts

The recent surge of interest in scRNA-seq stems from its ability to elucidate the correlation between genetics and diseases at the level of individual cell types with greater granularity [37–39]. To discern the cell-specific gene expression landscapes, numerous scRNA-seq analysis tools have emerged. The reliability of these tools in practical applications hinges on their evaluation against realistically labelled data [40,41]. Consequently, many simulators have been developed to ensure the availability of ground truth data [42–44]. Most of these simulations are based on statistical inference techniques, using well-estimated parameters of probability distributions to characterise given scRNA-seq data [45,46]. However, statistical inference methods have two limitations: long training time and a lack of denoising strategies.

Firstly, methods in this category typically lack technical support from computing acceleration technologies, such as graphics processing units (GPUs), particularly when compared to deep learning computing libraries. However, the practical utility of simulated data varies across cell, tissue, and disease types, potentially necessitating model retraining to accommodate a range of application conditions. On the other hand, the inherent variability of complex biological systems often eludes capture by rigid statistical inference methods that rely solely on fixed probability distribution assumptions [47,48]. This limitation may constrain their ability to effectively denoise data exhibiting multimodal distributions.

To address these limitations, we utilised a variant of the variational autoencoder (VAE) [49], the conditional VAE (CVAE) [50,51], to simulate transcriptomics data conditioned on cell types. The CVAE allows for training simulators on highly dimensional, yet sparse single-cell gene expression data at a significantly accelerated speed. Additionally, it integrates various types of neural networks into the parameterized statistical inference process, offering greater flexibility than purely statistical inference methods for denoising multimodally distributed data.

To train a CVAE-based single-cell gene expression simulator, we construct a log-likelihood term 𝓁 (*θ, φ*,**x**,**y**) as the optimisation objective

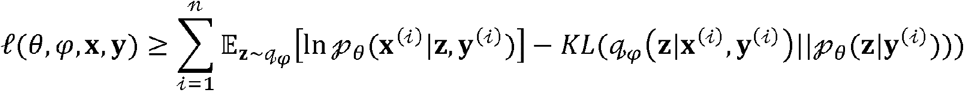

where 𝓆, is the encoder (the inference model) paramaterised *φ* to construct the statistical terms of a latent variable **z**, using the gene expression values of a cell .**x**^(*𝒾*)^ corresponding to type **y**^(*𝒾*)^. The decoder (the generative model) parameterized by *θ*, generates new single-cell data samples using the constructed **z**. *KL* represents the Kullback–Leibler (KL) divergence [52] quantifying the difference between the two distributions 𝓆and 𝓅 from their respective generative and inference processes. The full description of the modelling process can be found in section *Methods*.

The use of the latent variable **z** is popular in single-cell studies for addressing data denoising, perturbation, and batch effects. To better fit distributions 𝓆 and 𝓅, we employed convolutional neural networks (CNNs) [53] in both the encoder and the decoder due to their reduced parameters and fast training speed compared to fully connected neural networks [54] (**Figure 6**). However, CNNs can be replaced by any other types of neural networks, providing flexibility for users.

**Figure 6.**
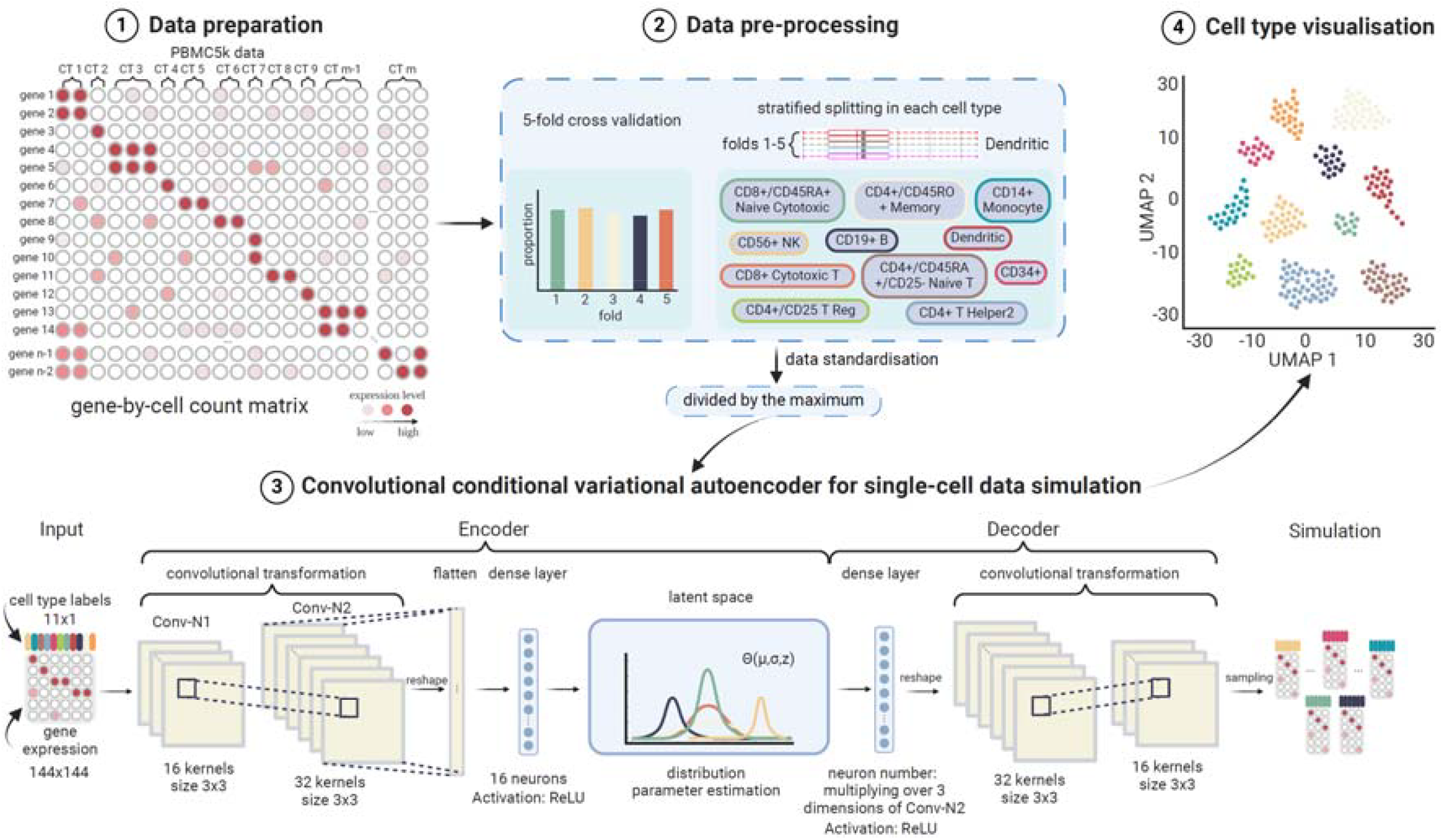
DeepConvCVAE for UMI count matrix simulation. It includes data preparation, data pre-processing, learning process, and cell type visualisation. 68k PBMCs scRNA-seq data was used and downloaded from https://github.com/10XGenomics/single-cell-3prime-paper/tree/master/pbmc68k_analysis. Single-cell data are filtered without a cell expressing below three genes. We used a stratified 5-fold cross validation to split the datasets for training. This ensures the cells from each cell type are mostly even across the 5 folds. Single-cell data were transformed through convolutional neural networks (CNNs) in the encoder and decoder of the conditional variational autoencoder. To suit the input size to the CNN, the single-cell count matrix is reshaped as a 144×144 square matrix. The same settings were used in CNNs in both the encoder and the decoder.

The trained model simulated gene expression data of 2000 cells for each of 11 cell clusters (**Figure 7**). After 10 training epochs, the simulated data has been shown discriminative between different cell clusters observed in PCA [55], TSNE [56], and UMAP [57] plots. We also integrated the overall framework of DeepConvCVAE into UMIche, for data simulation using trained models, data retraining, and framework modifications for other single-cell objective-specific studies.

**Figure 7.**
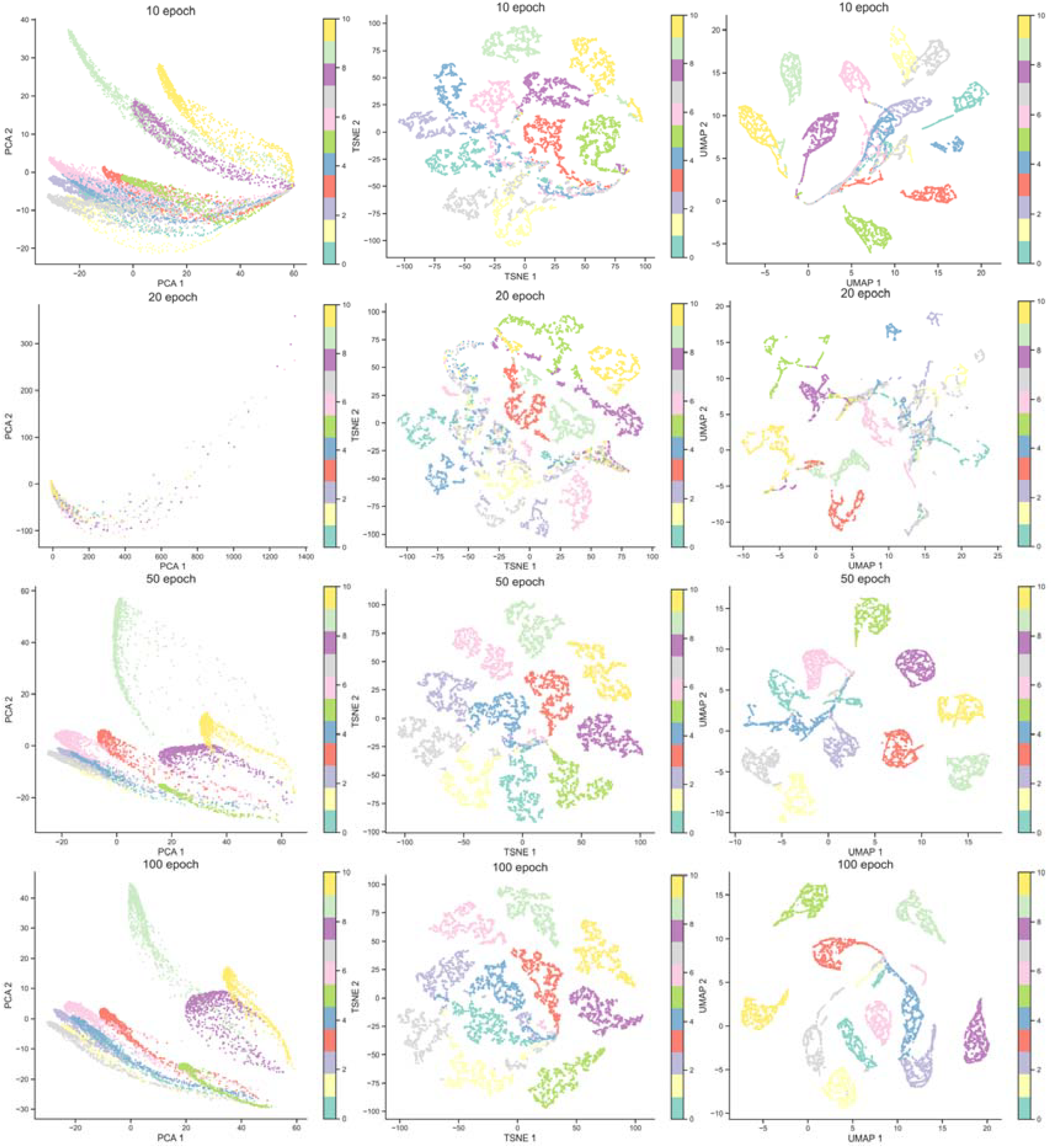
UMAP plots of simulated UMI counts by DeepConvCVAE at 10, 20, 50, and 100 epochs.

## Conclusions

In this work, we introduced UMIche, a comprehensive platform specifically designed for UMI-centric analysis, supporting the development and validation of UMI collapsing methods using both simulated and experimental bulk RNA-seq or scRNA-seq data. It offers four main modules: UMI collapsing method implementation, UMI collapsing pipeline implementation, UMI count matrix implementation, and visualization of UMI metrics. These modules are implemented to allow easy integration into existing computational programs and analysis pipelines. We present the architecture and functionalities of UMIche, along with case studies demonstrating its utility in advancing UMI deduplication performance in single-cell sequencing studies. UMIche has important implications for generating novel methodological insights by integrating recent advances in utilizing homotrimer UMIs for deduplicating PCR artefacts. This integration leverages error-tolerant recurrence patterns, offering a promising approach to enhance UMI collapsing methods and safeguard the accessibility to highly accurate gene expression profiles. We expect the use of UMIche to bridge measuring genomic observations and understanding biological mechanisms.

## Method

### UMI collapsing

UMI collapsing plays an important role in error correction and molecular quantification after sequencing [27]. The development of homotrimer UMIs makes it possible to collapse within UMIs before/after collapsing between homotrimer UMIs [31]. However, it remains murky as to how much we can gain in improving UMI collapsing effects from their combination. In UMIche, we constructed a homotrimer UMI-based collapsing scheme by combining both intra-UMI collapsing and inter-UMI collapsing methods, which is formulated below.

### Step 1: Splitting homotrimer UMIs into monomer UMIs

Let 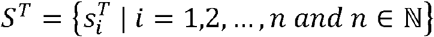 be the set of *n* homotrimer UMIs of length *l* observed at a single genomic locus. Let *W=*{*A,T,C,G*} be the set of four types of nucleotides. For any homotrimer UMI 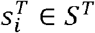, its *j* th trimer block is represented by set 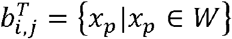 for which 𝓅 ∈ ={1,2,3} is a set of positional coordinates of elements in 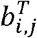. The set of all of its trimer blocks 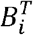 is denoted as

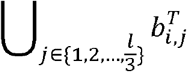

Consider function 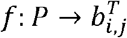 for projecting a positional coordinate into a nucleotide contained in 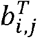. We aim to replace a trimer block 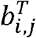 with a new set 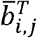, in which heterogeneity among nucleotides is obviated by the majority vote strategy. We retain 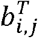 for 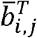 if given 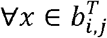, ∄𝓅_1_ ≠ 𝓅_2_, 𝓅_1_ ∈*P* and 𝓅_2_ ∈*P*, such that *f* (𝓅_1_)= *f* (𝓅_*2*_)=*x*. Otherwise, we construct 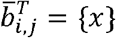. Accordingly, 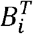 turns to 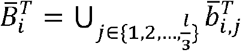.

Let *F* be a function for splitting a homotrimer UMI into monomer UMIs. Specifically, it uses as input the set of trimer blocks of a homotrimer UMI, takes turn selecting a nucleotide from each homotrimer block, and finally merges those selected nucleotides into monomer UMIs.

According to whether or not to apply *f* to each trimer block, utilising *F* to obtain split monomer UMIs results in two respective sets of sequences.

*spALL*. If *f* is not applied, we obtained a set containing all possible monomer UMIs from a homotrimer UMI. We term this strategy *split-to-all* (*spALL*). The resulting set of its split *n*_*i*_ monomer UMIs is phased to be

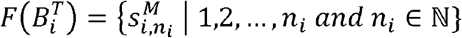

*spMV*. Let 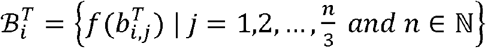 denote the set of heterogeneity-obviated trimer blocks of a homotrimer UMI 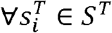. Thus, its split *m*_*i*_ monomer UMIs comprise the following set

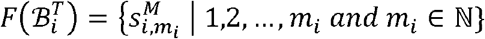

We term this strategy *split-by-majority-vote* (*spMV*).

### Step 2: Homotrimer UMI deduplication (deduplication stage 1)

#### 1. Deduplication using set cover optimisation

Let *g* be the set cover optimisation function [58,59] for removing PCR artefacts using a set of homotrimer UMIs at a given genomic locus. The deduplicated count *c* at the locus is written as

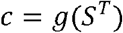

To solve function *g*, we use the greedy algorithm [33], which iteratively finds subsets of homotrimer UMIs from set *S*^*T*^ through their overlapped monomer UMIs with the maximum counts. At each iteration, specifically, the greedy algorithm attempts to identify a monomer UMI *s*^*M*^ that can account for a subset of the largest number of homotrimer UMIs. The homotrimer UMIs are thought to derive from the same original homotrimer UMI at the locus, which are then popped out of *S*^*T*^ in the end of each iteration. The remaining homotrimer UMIs in *S*^*T*^ are iteratively deduplicated likewise, until none of monomer UMIs is found to account for more than one homotrimer UMI. Suppose that there are *k* iterations to identify these *k* monomer UMIs, each accounting for at least two homotrimer UMIs. There are *r* monomer UMIs left, each corresponding to only one homotrimer UMI. Hence, the final deduplicated count *c* is the sum of the both, like that,

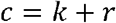

This implies that the problem of identifying PCR duplicates at the level of homotrimer UMIs is converted to the problem of settling the optimised coverage between sets at the level of monomer UMIs.

Let *S*^*K*^ ={*n*_*i*_ ∣*n*_*i*_ ∈ ℕ *for i=*0.1,2, …,*k*} be the set of the numbers of homotrimer UMIs in *k* subsets 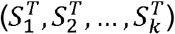 for which monomer UMIs account after *k* iterations. Note that *n*_0_ = 0. In each iteration, the identified homotrimer UMIs are collected into 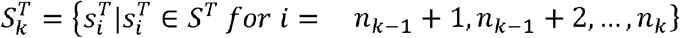. The union of the *k* subsets is a collection of all homotrimer UMIs in which at least two can be accounted for by one monomer UMI, denoted as

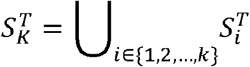

Let 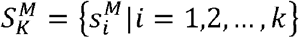 be the set of these monomer UMIs after *k* iterations. The remaining homotrimer UMIs, whose split monomer UMIs are unable to be overlapped with other homotrimer UMIs, are stored in set 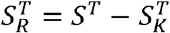. Let 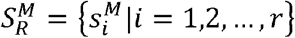 be the set of these monomer UMIs after *k* iterations.

##### Definition 1.1.

Let 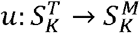 be a **many-to-one** mapping from homotrimer UMIs to monomer UMIs. Original homotrimer UMIs are considered to have duplicates if given 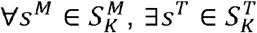, such that

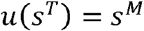

##### Definition 1.2.

Let 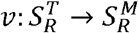 be a **one-to-one** mapping from homotrimer UMIs to monomer UMIs. Original homotrimer UMIs are not considered to have duplicates if given 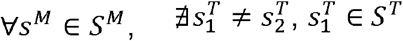 and 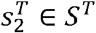, such that

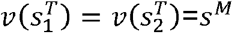

#### 2. Deduplication using majority voting

Let *h* be the majority voting function for removing PCR artefacts using a set of homotrimer UMIs at a given genomic locus. The deduplicated count at the locus is written as

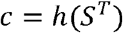

To solve function *h*, we begin by obtaining a sole monomer UMI that corresponds to a homotrimer UMI. This is achieved by collapsing a trimer block 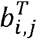 into a single nucleotide by the majority vote strategy and then concatenating all single nucleotides into a monomer UMI. S. We collapse 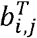 into the recurring nucleotide *x* if given 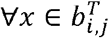, ∃ 𝓅_1_ ≠ 𝓅_2_. 𝓅_1_ ∈*P* and 𝓅_2_ ∈*P*, such that *f*(*p*_1_)*= f*(*p*_2_)*=x*. Otherwise, a random sampling operation is applied to obtain a nucleotide 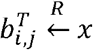. Suppose that 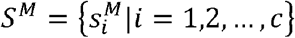 is the set of unrepeated monomer UMIs collapsed from homotrimer UMIs in *S*^*T*^. In this case, the number of the unrepeated monomer UMIs is said to be the deduplicated count *c*.

Let *S*^*C*^ ={*n*_*i*_ ∣*n*_*i*_ ∈ ℕ *for i=*0.1,2, …,*c*} be the set of the numbers of homotrimer UMIs in *c*. subsets 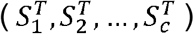. Note that *n*_0_ =0. Each subset 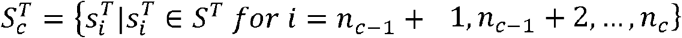 contains homotrimer UMIs that are all collapsed into a common monomer UMI. The union of the *c* subsets is set *S*^*T*^, denoted as

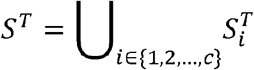

##### Definition 2.1.

Let *w*: *S*^*T*^ → *S*^*M*^ be a **many-to-one** mapping from homotrimer UMIs to monomer UMIs. Original homotrimer UMIs are considered to have duplicates if given ∀*s*^*M*^ ∈ *S*^*M*^, ∃ *s*^*T*^ ∈ *S*^*T*^, such that

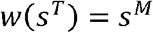

### Step 3: Retaining homotrimer UMIs

#### 1. Set cover optimisation

We randomly sample homotrimer UMIs from the *k* subsets 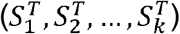, respectively. This forms a new set 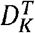, given by

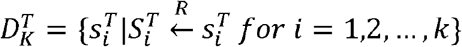

Then, together with set 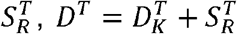 is taken as the set that comprises homotrimer UMIs that are one-to-one mapped to all identified monomer UMIs.

#### 2. Majority voting

We randomly sample homotrimer UMIs from the *c* subsets 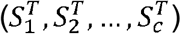 respectively, leading to

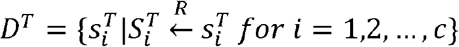

*D*^*T*^ is said to be the set that comprises homotrimer UMIs that are one-to-one mapped to all monomer UMIs collapsed by the majority vote strategy.

### Step 4: Collapsing homotrimer UMI

#### 1. Collapsing blocks by the majority vote strategy (*cbMV*)

We define function 𝓆to concatenate a set of nucleotides into a sequence. Let 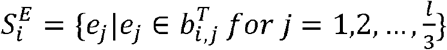 be a set of nucleotides, each added by collapsing a trimer block 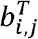 of any homotrimer UMI 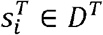 into a single nucleotide by the majority vote strategy. *e*_*j*_ is nucleotide *x* if given 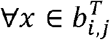 ∃ 𝓅_1_ ≠ 𝓅_2_. 𝓅_1_ ∈*P* and 𝓅_2_ ∈*P*, such that *f*(*p*_1_)*= f*(*p*_2_)*=x*. Otherwise, a random sampling operation is applied, such that 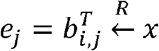. Subsequently, we build set 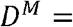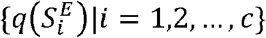, which adds each concatenated monomer UMI for each homotrimer UMI 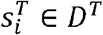.

#### 2. Collapsing blocks randomly (*cbRAN*)

We define function to concatenate a set of nucleotides into a sequence. Let 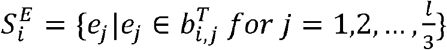 be a set of nucleotides, each added by collapsing a trimer block 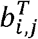 of any homotrimer UMI 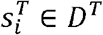 into a single nucleotide through the random sampling strategy, such that 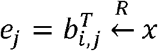. Then, we build set 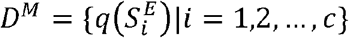, which adds each concatenated monomer UMI for each homotrimer UMI 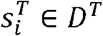.

### Step 5: Homotrimer UMI deduplication (deduplication stage 2)

Through steps 1-4, a collection of homotrimer UMIs in *S*^*T*^ are deduplicated into a bundle of monomer UMIs in *D*^*T*^ Finally, an external UMI deduplication method *φ*, which relies typically on the use of graph modelling techniques, is applied on *D*^*T*^ to further remove UMI deduplicates. The calculation result *φ*(*D*^*T*^) is deemed as the deduplication count *c* for this step.

### UMI count matrix simulation

Let **x** be a count matrix with 𝓃 cells and 𝓂 genes in the following form.

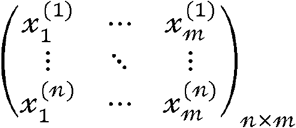

Cell types of the 𝓃 cells are known and stored in vector **y** = [𝓎_1_, 𝓎_2,_ … 𝓎_*n*_] in which there are 𝓁 unique cell types in total. In 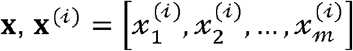 represents expression values of 𝓂 genes for cell 𝒾 whose cell type corresponds to 𝓎 _𝒾_ in **y**. We might as well consider fitting **x**, to a probability distribution function 𝓅 first and then drawing data from this distribution to approximate **x** as the use of probabilistic models is usually thought of to be more generalizable and preferable to capture and discover the underlying patterns of biological systems and gene regulation mechanisms behind the variable, yet incompletely characterized biological data, such as single-cell count matrices. The goal of the probabilistic modeling process is set to estimate parameters *θ* from *𝓅* so as to approximate **x**, which is described as

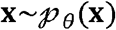

As the effect of approximation of *θ* may be affected by unknown patterns that are unobservable from **x**, a latent variable **z** needs to be introduced to settle this problem [60], which turns 𝓅_θ_ (**x**) to a conditional distribution 𝓅_θ_ (**x**| **z**) from which **x** is about to be drawn. Consider the following probabilistic model [61],

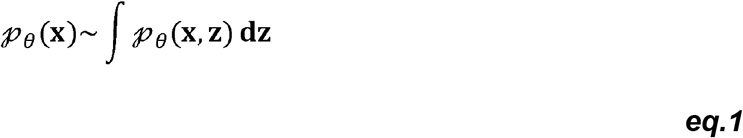

According to Bayes’ theorem 𝓅_θ_ (**x, z**) =𝓅_θ_ (**x**| **z**) 𝓅_θ_ (**z**)= 𝓅_θ_ (**z**| **x**) 𝓅_θ_ (**x**) it turns to

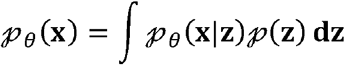

Due to the intractability to solve the above integral [49,62], we are unable to obtain the suitable estimates of 𝓅_θ_ (**x**| **z**) As for the prior distribution of 𝓅 (**z**), it is more practical to consider its posterior distribution 𝓅(**z**| **x**)as it is conditioned on the observations **x**, i.e., the realistic posterior distribution after the model in ***eq*.*1*** is fed with **x**. However, after the formula transformation below, it is also intractable to solve 𝓅_θ_ (**z**| **x**) according to [63].

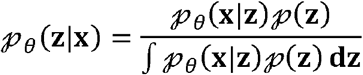

Hence, we sought to find another distribution 𝓆 _φ_ (**z**| **x**) parameterized by φ to substitute 𝓅_θ_ (**x**| **z**), such that,

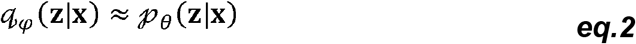

Reconsider the probabilistic model present in ***eq*.*1*** before solving 𝓆 _φ_ (**z**| **x**) .To estimate *θ* from the model, it is common to apply a log-likelihood function as follows

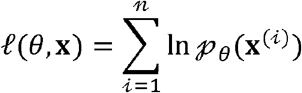

For a given single cell is written as

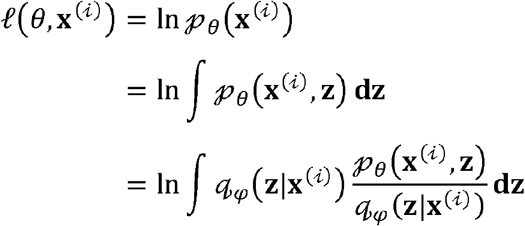

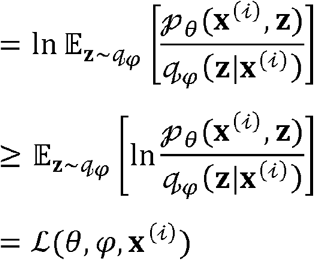

ℒ(θ, φ, **x**^(𝒾)^) is known as the evidence lower bound (ELBO) function of 𝓁 (θ **x**^(𝒾)^) [64]. After applied with Bayes’ theorem, ℒ is further written to be the following form

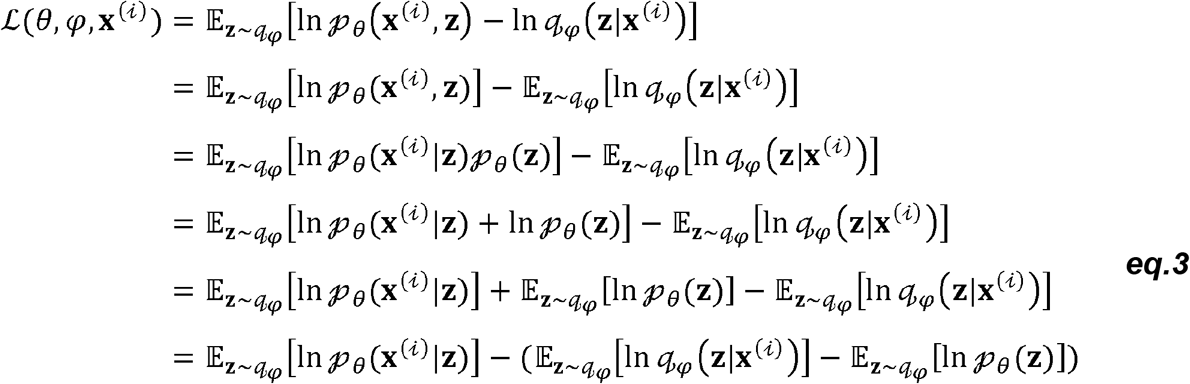

Thus, altogether, the log-likelihoods of all observed gene expression values across 𝓃 single cells with respect to its ELBO (the right side of the equation) are expressed as

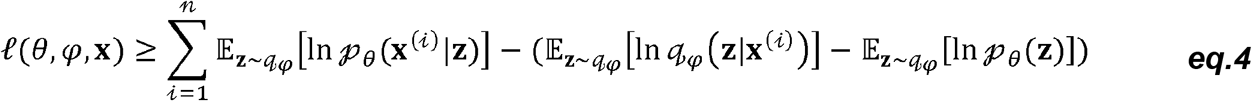

Let ℒ_1_ be 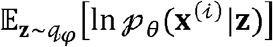, which is a term that we expect to use to reconstruct data based on θ-parameterized distribution 𝓅 conditioned on **z**. Let ℒ_2_ 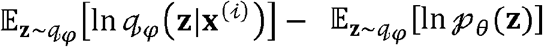. In fact, ℒ_2_ is the Kullback–Leibler (KL) divergence that measures the discrepancy between two distributions [52]. Recall ***eq*.*2***, compared with the estimation of prior distributions of **z** it is likely more rational to estimate its posterior distributions if given the observed data x^(𝒾)^. Hence, ℒ_2_ is reformatted as

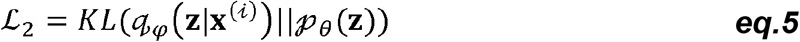

According to ***eq*.*5***, it becomes clear that a perfect estimation of the true posterior distribution 𝓅_θ_ (**z**| **x**^(𝒾)^) can be achieved if ℒ_2_ is 0, suggesting that a substituted distribution of 𝓆 _φ_ (**z**| **x**^(𝒾)^) bears a very close resemblance to 𝓅_θ_ (**z**).

ℒ_1_ is often referred to as the generative model (**the decoder**) to reconstruct **x**, while ℒ_2_ is often referred to as the inference model (**the encoder**) to build a Gaussian distribution of input **x**. An autoencoder with such a setting, which learns a distribution from input data and use the learned distribution to simulate new data, is referred to as the variational autoencoder (VAE) [49].

Let 𝓅_θ_ (**z**).be a multivariate Gaussian distribution 𝒩 (0,I) where I represents a standard diagonal covariance matrix. Let 𝓆 _*φ*_ (**z**| **x**^(𝒾)^) be a multivariate Gaussian distribution 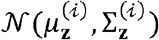 with respect to the gene expression values of cell 𝒾. 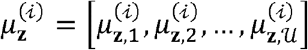 is a vector that shares the same dimension of 𝒰 to the latent variable **z**. 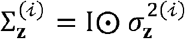 (⊙ for element-wise multiplication) is a diagonal covariance matrix whose diagonal elements are 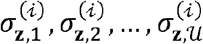. Then, combined with the multivariate Gaussian function, ***eq*.*5*** is finally reformatted as

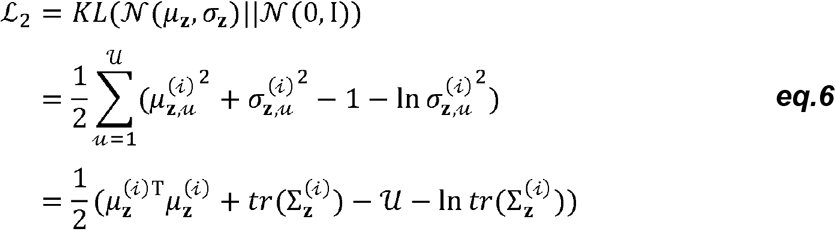

According to ***eq*.*6***, the inference model in the encoder is set up for learning two Gaussian distribution parameters, *μ*_**z**_ and Σ_**z**_, from the single-cell count matrix **x**. In our case, to perform mapping 𝓆 _*φ*_ (**z**| **x**^(𝒾)^): **x** → *μ*_**z**_, Σ_**z**_, we employed a convolutional neural network [54] and a multiple perceptron (with 1 dense layer with 16 neurons) [65] to transform input**x** into values ranging from 0 to 1 to approximate *μ*_**z**_ and Σ_**z**_. The convolutional neural network comprises two layers, with the first placed with 16 kernels with a size of 3 ×3 and the second placed with 32 kernels with a size of 3 ×3 (**Figure 6**).

Before performing the reconstruction of single-cell gene expression data approximate to. in the decoder, we made use of the output of the encoder, *μ*_**z**_ and Σ_**z**_, to build **z** taken as input to the decoder. Following the common practice, we employed the reparameterization trick, calculated by

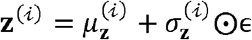

where ϵ is a 𝒰-dimension vector whose elements are drawn from another Gaussian distribution 𝒩 (0,I). In our case, to avoid very small values during training, we artificially added a value 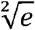 to 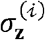. Then, **z**^(𝒾)^ is transformed to

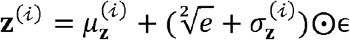

In the decoder, we placed the same neural network architecture with the encoder (**Figure 6**). As the decoder requires the reconstructed **x** to be as more similar as possible, the loss of the decoder is set to minimise the cross entropy (*CE*) [66] between the ground-truth and simulated gene expression values. The loss of the encoder is ℒ_2_ itself (***eq*.*6***) as the KL divergence aims to contract the difference between a posterior distribution of **z** under 𝓆 _*φ*_, and its prior distribution under 𝓅 _θ_. Thus, the total loss of the model is the joint loss of the encoder and the decoder, given by

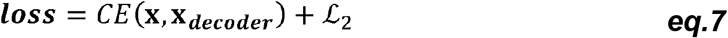

We used the root mean square propagation technique (RMSProp) [67] to optimize the model and set the batch size to 32 cells. Using convolutional neural networks as both the encoder and the decoder is the significant reduction of parameters required to train and simulate the single-cell gene expression data. Although the above model is capable of generating gene expression data, it is still unable to produce cell type-specific gene expression data. Similar to introducing the latent variable that observable data **x** is conditioned on, **y** can be seen as another condition further introduced to the model in ***eq*.*4***, termed CVAE [50,51] and rewritten as

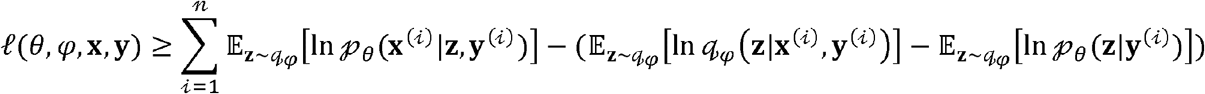

To this end, a simple operation further required in coding is the concatenation of **x** and **y** as the input to both the encoder and the decoder. As a result, the cell types specific to their relative gene expression data are together trained and learned by the VAE model.

### Training the VAE model on the 68k PBMC dataset

We pulled from 10xGenomics the *68k PBMCs Donor A* dataset with 32,738 genes expressed in 68,579 cells [68]. Each cell has at least one expressed gene. We left with 20,387 genes after removing genes with zero expressions in all cells. Next, genes that are expressed in less than 20 cells are also removed. The cell clusters were detected from barcodes using the CellRanger 1.1 pipeline and obtained under tab “*Zheng et al*.” from https://github.com/10XGenomics/single-cell-3prime-paper/tree/master/pbmc68k_analysis. Then, the processed dataset was split into 5 folds using the stratified shuffle split method to ensure that cells from different cell clusters (CD8+ Cytotoxic T, CD8+/CD45RA+ Naive Cytotoxic, CD4+/CD45RO+ Memory, CD19+ B, CD4+/CD25 T Reg, CD56+ NK, CD4+ T Helper2, CD4+/CD45RA+/CD25-Naive T, CD34+, Dendritic, and CD14+ Monocyte) were evenly allocated in each fold (**Supplementary Figs. 6-8**). Data were normalized into 0-1 prior to training. The parameters of the final model were optimized on each training fold. All operations above were automatically performed with Tensorflow [69] and Keras [70], the trained model DeepConvCVAE is made available in UMIche.

## Data and code availability

The standalone package of UMIche is made available at https://github.com/cribbslab/umiche. The data used in this paper can be accessed through https://cribbslab.github.io/umiche. The *68k PBMCs Donor A* dataset can be downloaded through tab “*Zheng et al*.” at https://www.10xgenomics.com/resources/datasets and the information about their cell types can be found at https://github.com/10XGenomics/single-cell-3prime-paper/tree/master/pbmc68k_analysis.

## Declaration of competing interest

A.P.C is listed as an inventor on several patents filed by Oxford University Innovations concerning single-cell sequencing technologies. The other authors declare that they have no known competing financial interests or personal relationships that could have appeared to influence the work reported in this paper.

## Author contribution

J.S. conceived this research. J.S. performed the data analysis, plotted the data, developed the software, and wrote the manuscript. J.S., S.L., and S.C. contributed to the development of the set coverage optimisation method and computational analyses. J.S. and A.P.C. revised the manuscript and acquired funding. All authors have given approval to the final version of the manuscript.

## Acknowledgement

This work was financially supported by the Medical Research Council (MRC) career development fellowship (MR/V010182/1).

## Notes

### Summary of Updates

The corresponding authors were not correctly designated in the initial version. We have submitted a revised version to rectify this.

