## Supplementary Information for "UMIche: A platform for robust UMI-centric simulation and analysis in bulk and single-cell sequencing"

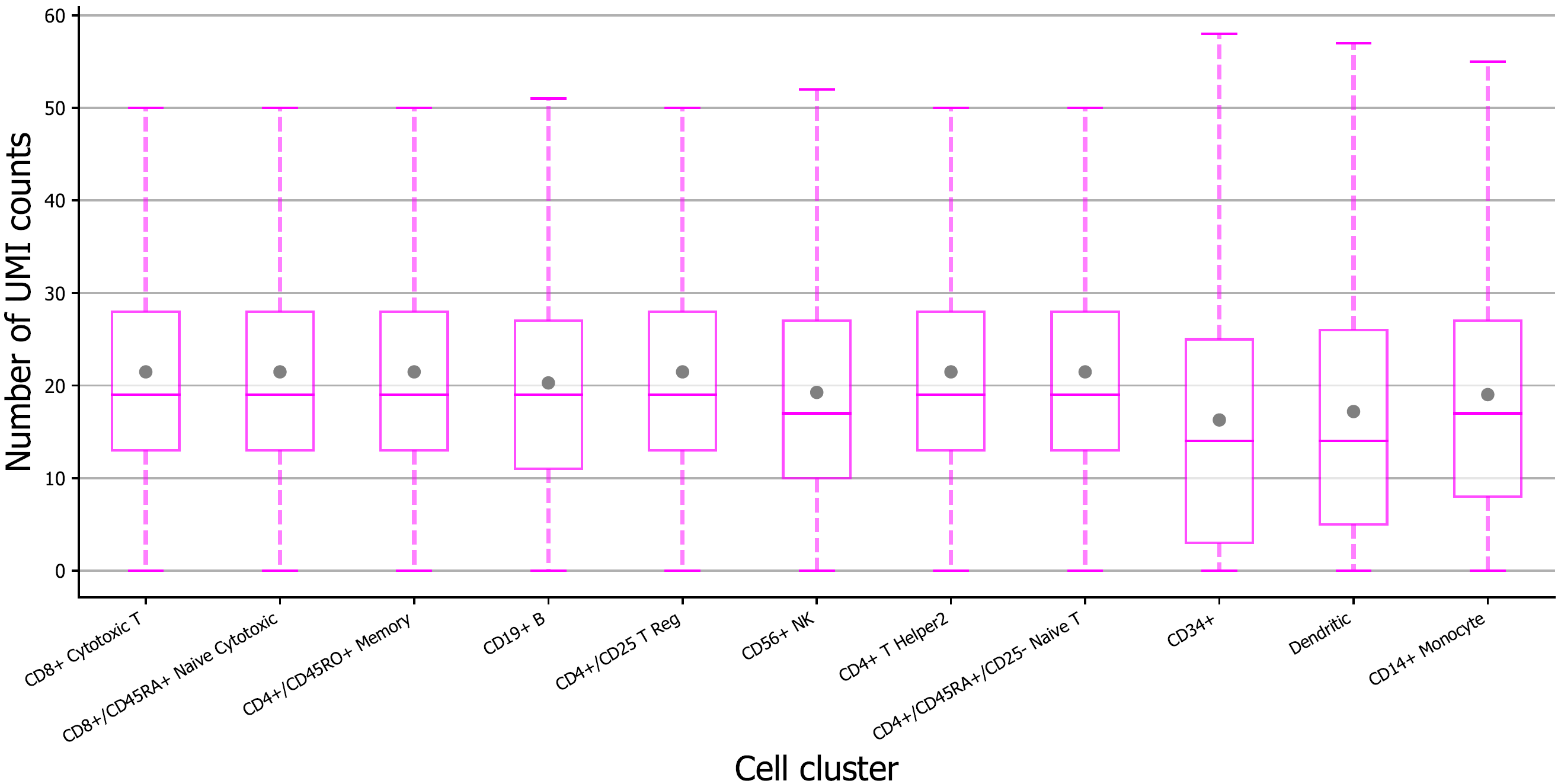

**Supplementary Figure 1.** UMI counts of top 10 highly expressed genes in the 68k PBMCs Donor A dataset at per cell cluster level.

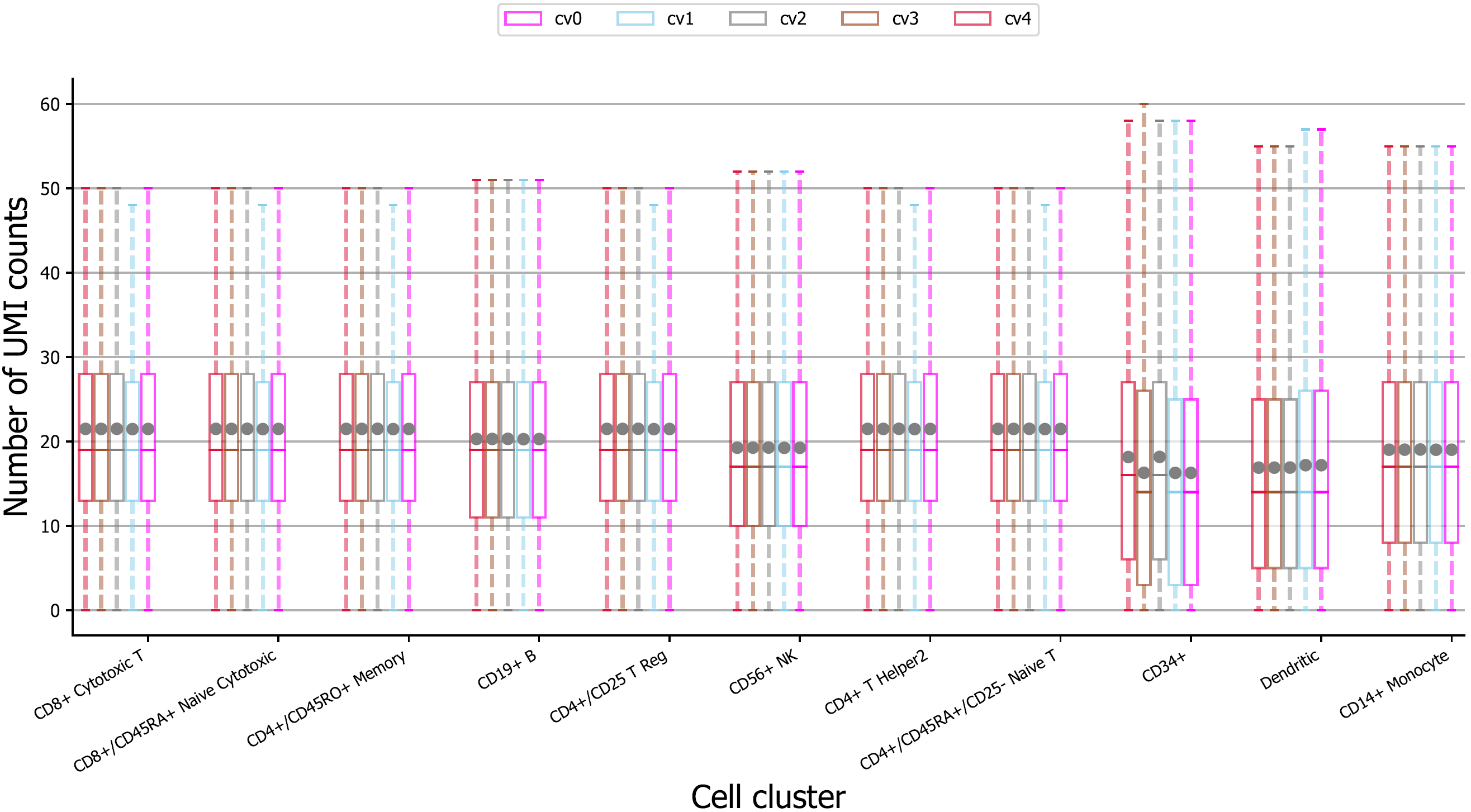

**Supplementary Figure 2.** UMI counts of top 10 highly expressed genes in 5-fold cross validation (cv) sets for training, including cv0, cv1, cv2, cv3, and cv4, spilt by the stratified shuffle split method at per cell cluster level.

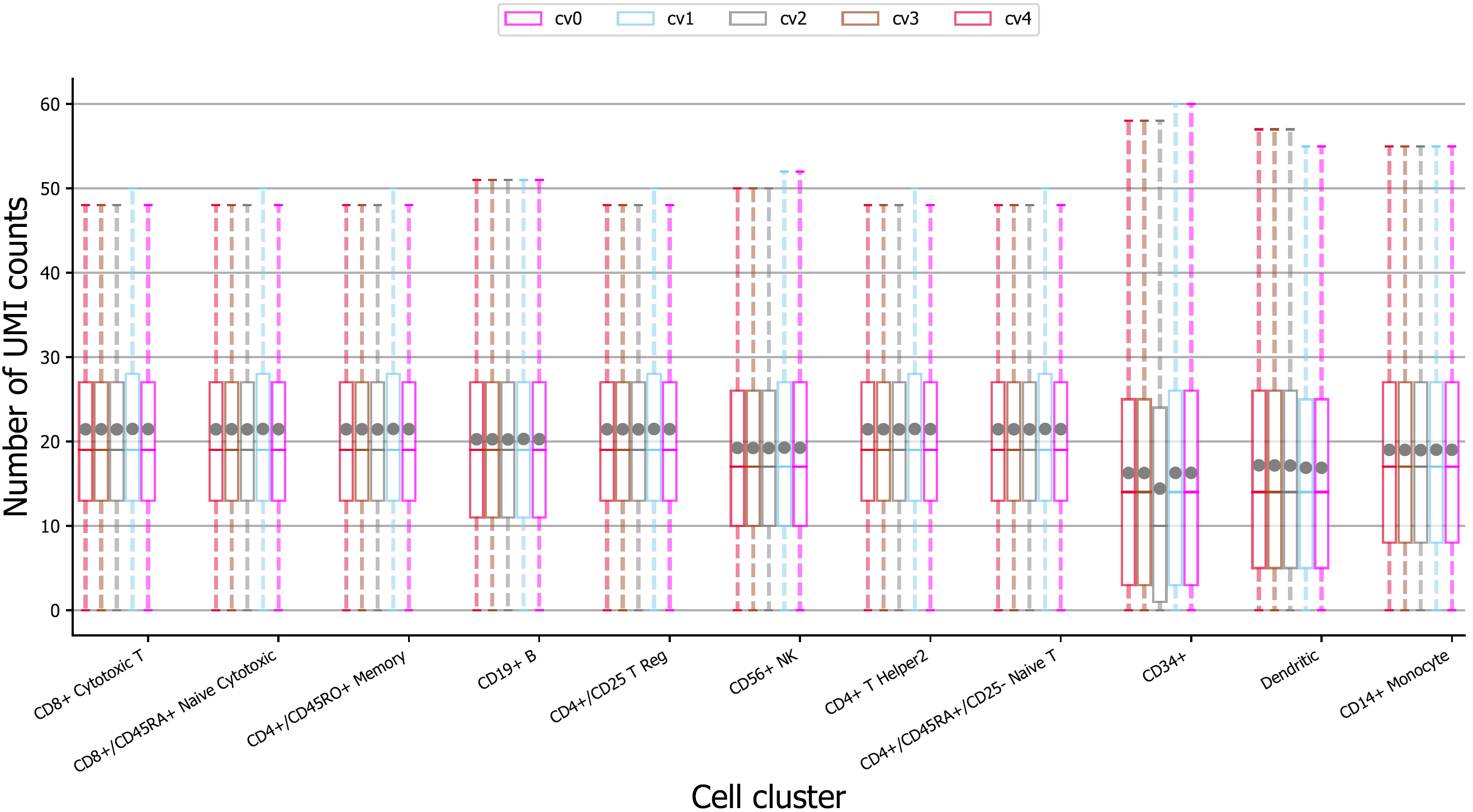

**Supplementary Figure 3.** UMI counts of top 10 highly expressed genes in 5-fold cross validation (cv) sets for validation, including cv0, cv1, cv2, cv3, and cv4, spilt by the stratified shuffle split method at per cell cluster level.
